# Phosphorylation alters the bulk chemical properties of Orc1 to tune DNA binding, phase separation, and heterochromatin partitioning

**DOI:** 10.64898/2026.08.17.745304

**Authors:** Olubu A. Adiji, Innesa Leonovich, Matthew W. Parker

## Abstract

The first step in initiating DNA replication is binding of the origin recognition complex (ORC) to chromosomes. Metazoan ORC is recruited to chromatin via the Orc1 intrinsically disordered region (IDR) whose DNA and chromatin binding activity are regulated by Cyclin Dependent Kinase (CDK) phosphorylation. ORC is also enriched in heterochromatin where it is required for the formation and maintenance of a silenced chromatin state. ORC’s recruitment to heterochromatin is developmentally and cell cycle regulated but the underlying regulatory mechanism remains unknown. We hypothesized that CDK-dependent phosphorylation of the Orc1 IDR underpins regulated recruitment to heterochromatin. Using bioinformatic analyses, we find that the *Drosophila* Orc1 IDR (Orc1^IDR^) contains an exceptionally high density of CDK phospho-sites and, despite considerable sequence variation, the density of sites, but not their position, is conserved. *In vitro* DNA binding and phase separation experiments reveal that phosphorylation tunes Orc1^IDR^ function in a rheostat-like fashion. Using phospho-mimetic variants, we find that constitutive phosphorylation not only weakens interphase chromatin binding but fully inhibits partitioning of Orc1^IDR^ into heterochromatin. Finally, we use phospho-mimetic variants to probe the importance of site-specific phosphorylation and find that the precise position of sites can be changed provided the new sites are equitably distributed across the sequence. These studies demonstrate that phosphorylation tunes the biochemical properties of the Orc1 IDR to control DNA binding, phase separation, and, consequentially, heterochromatin recruitment. This work suggests that localized dephosphorylation of the DNA binding Orc1 IDR may underlie recruitment of ORC to specific genomic loci.

## INTRODUCTION

DNA replication is a fundamental process of life and serves to maintain genome stability through cycles of cell division. Preparation for eukaryotic DNA replication begins in late mitosis and early G1 phase when chromatin is licensed for replication by loading of the Mcm2-7 replicative helicase around duplex DNA by three factors: the Origin Recognition Complex (ORC), composed of Orc1-6 subunits, Cdc6, and Cdt1 (reviewed in (1)). In addition to their role in normal cell physiology, replication licensing factors are commonly overexpressed in cancer where they are implicated in driving genome instability (2) and germline mutations in the same factors are known to cause Meier-Gorlin Syndrome (MGS), a form of primordial dwarfism (3).

The first step of DNA replication licensing is binding of ORC to specific genomic loci, known as origins, where Mcm2-7 loading occurs (4, 5). The mechanism of origin selection is well-understood in budding and fission yeast with each ORC ortholog possessing a species-specific auxiliary DNA binding appendage that mediates binding to specific DNA sequences (6–8). Conversely, the mechanism of origin selection in metazoans has remained elusive. Our recent work demonstrates that Orc1 contains an intrinsically disordered region (IDR) that binds DNA non-specifically *in vitro* and which is necessary and sufficient for chromatin recruitment of ORC in the early *Drosophila* embryo (9, 10). Whether the Orc1 IDR also contributes to loci-specific chromosomal binding is currently unknown. Importantly, phosphorylation of the Orc1 IDR by Cyclin Dependent Kinases (CDKs) inhibits its DNA and chromatin binding activity (10) and thereby restricts ORC’s replication licensing function to late mitosis and early G1 phase of the cell cycle when CDK activity is low (11).

Beyond its role in replication, yeast and metazoan ORC also play a role in transcriptional silencing (12–14). Metazoan ORC is required for the maintenance of heterochromatin and loss of ORC results in a position-effect variegation (PEV) phenotype in flies (14) and physical reorganization of heterochromatin in human cells (15, 16). Human ORC is labile (17–19) and the Orc1 and Orc2-5 sub-complexes interact with heterochromatin proteins Hp1*α* (14, 16) and ORC-associated protein (ORCA) (20, 21), respectively. Interestingly, the formation of heterochromatin is now understood to be intimately linked to the physical properties of resident factors, with both Hp1*α* and ORC able to mediate DNA-dependent phase separation *in vitro*, which has been hypothesized to underlie the formation of a liquid-like condensed chromatin state (9, 22–25). Conversely, ORCA limits nucleosome array compaction which may make heterochromatin more suitable for licensing (26). Although recruitment of ORC to heterochromatin is developmentally (9) and cell cycle regulated (15, 16), how this regulation occurs is currently unknown. Notably, the phase separation propensity of ORC is negatively regulated by phosphorylation (9), providing a possible mechanism to regulate partitioning into heterochromatin.

Here we test the hypothesis that phosphorylation of the Orc1 IDR functions as a key cellular mechanism to regulate heterochromatin partitioning of Orc1. We find that despite poor sequence conservation, Orc1 IDR orthologs have an exceptionally high and conserved density of CDK phosphorylation sites. Biochemical reconstitutions show that progressive phosphorylation tunes the DNA binding affinity and phase separation propensity of the Orc1 IDR. Using phospho-mimetic variants, we demonstrate that phosphorylation fully ablates Orc1 partitioning into heterochromatin and that this is independent of any specific phosphorylation site, relying instead on changes to the overall bulk chemical properties of the sequence. These studies provide a functional mechanism for the regulated recruitment of ORC to heterochromatin and highlight the ability of the Orc1 IDR to facilitate site-specific chromatin binding.

## RESULTS

### The Orc1 IDR has an exceptionally high and conserved density of CDK phosphorylation sites

*Drosophila* Orc1 has a multi-domain architecture with an intrinsically disordered region (IDR) (**Fig. 1A**) that we have previously shown is necessary and sufficient for DNA and chromatin binding and DNA dependent phase separation (9). Embedded within the IDR are fifteen CDK consensus motifs, including eight minimal motifs (‘[S/T]P’, **Fig. 1A**, black hash marks) and seven full motifs (‘[S/T]Px[R/K]’, **Fig. 1A**, red hash marks). To determine how Orc1’s CDK site density compares with other proteins, we assessed site density on a proteome-wide level. Specifically, we calculated the density of minimal and full CDK motifs in all disordered regions (IDRs > 50 amino acids) and compared this to the fly Orc1 IDR, which has a CDK site density of 4.1 minimal and 1.9 full motifs per 100 residues (**Fig. 1B**). We found that >75% of the 10,482 IDRs in the fly proteome have no full CDK consensus motifs and 99% of IDRs have a full site density < 1.75, placing Orc1 in the top 1% of sequences (**Fig. 1B**, red circles). Likewise, the density of minimal motifs (‘[S/T]P’) in the Orc1 IDR places it within the top 5% of all disordered regions (**Fig. 1B**, black circles).

**Figure 1:**
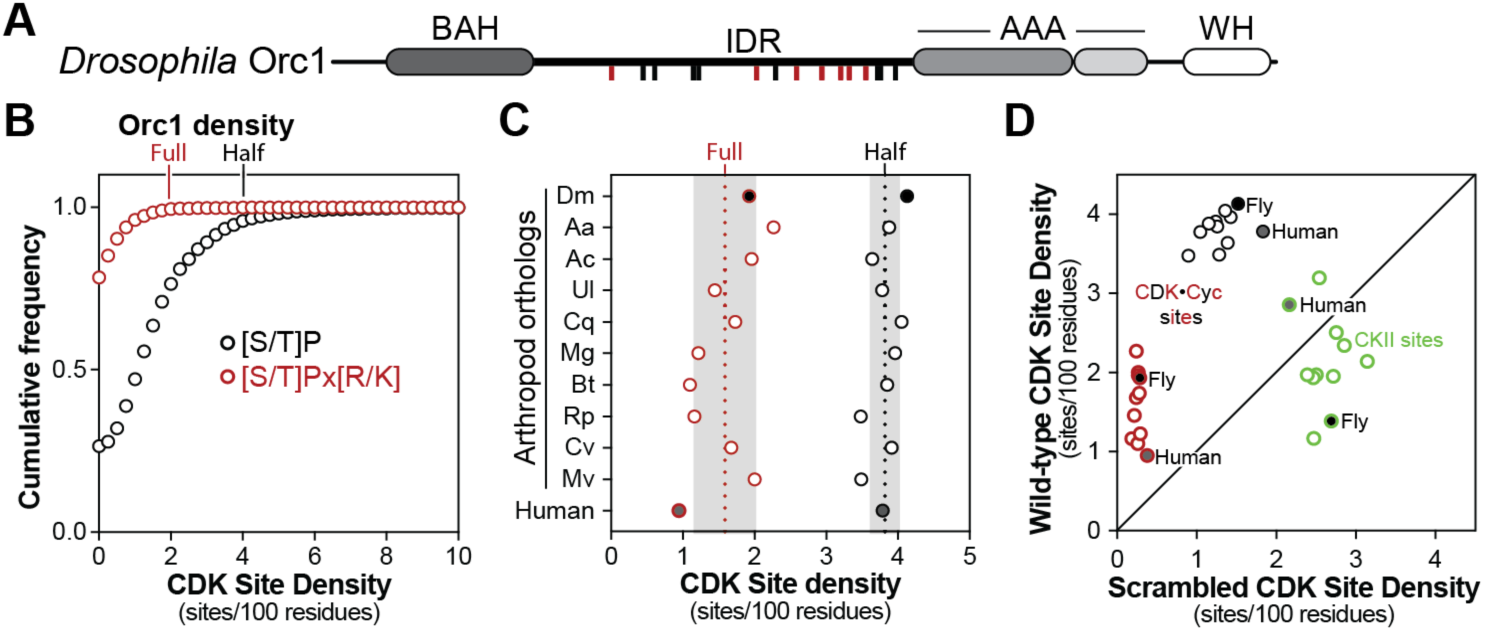
Analysis of CDK phospho-site density in the Orc1 IDR. A, architecture of fly Orc1. Red and black hash marks indicate full (‘[S/T]PX[R/K]’) or minimal (‘[S/T]P’) CDK phosphorylation motifs, respectively. Orc1 contains the following regions: a Bromo Adjacent Homology (BAH) domain, an intrinsically disordered region (IDR), an ATPase Associated with various cellular Activities (AAA) domain, and a winged-helix (WH) domain. B, the cumulative frequency of CDK site density in IDRs proteome wide. Data is shown for both full (red) and minimal (black) CDK sites. C, full (red) and minimal (black) CDK site density for arthropod and human Orc1 IDR orthologs. D, number of full (red) and minimal (black) CDK sites in human and arthropod Orc1 IDRs plotted against the average number of sites observed in 10,000 random scrambles of each sequence. The same analysis was done for CKII sites (green).

We next asked whether the unusually high density of phospho-sites is a conserved feature of Orc1 orthologs. We found that the density of both full (**Fig. 1C**, red circles) and minimal CDK sites (**Fig. 1C**, black circles) is maintained in nine arthropod and human Orc1 IDR orthologs. This conservation is notable given that the Orc1 IDR is highly divergent across the metazoan lineage such that human and *Drosophila* sequences cannot be aligned (10). To determine if CDK site density is simply a product of an amino acid sequence bias that renders these motifs more likely, we calculated the average density of minimal and full CDK consensus sites in 10,000 random scrambles of arthropod and human Orc1 IDR orthologs and plotted this against the number of observed sites. For all orthologs, the abundance of both full (**Fig. 1D**, red circles) and minimal CDK consensus sites (**Fig. 1D**, black circles) was higher than what was observed in randomly scrambled sequences. Conversely, the density of Casein Kinase II (CKII) phosphorylation motifs (‘[S/T]xx[D/E]’) was similar between Orc1 orthologs and their randomly scrambled counterparts (**Fig. 1D**, green circles). The conservation of CDK site density suggests that multi-site phosphorylation is critical for cellular control of Orc1 function.

### Progressive phosphorylation tunes Orc1 IDR DNA binding and phase separation

We reasoned that multi-site phosphorylation could regulate Orc1 IDR function through either a two-state, switch-like mechanism (i.e., on/off) or through tunable, rheostat-like control. These two models make separate predictions regarding the impact of partial versus full phosphorylation on Orc1 IDR function and we therefore prepared Orc1 IDR with varying levels of phosphorylation. Specifically, we treated the purified *D. melanogaster* Orc1 IDR (Orc1^IDR^) with recombinant CDK2•CycE and, by varying both the concentration of kinase and the time of the reaction, we were able to achieve differential phosphorylation. Each phosphorylated variant was subject to a final size exclusion chromatography step to remove CDK2•CycE and the final purified protein was assessed by SDS-PAGE (**Fig. 2A**, top) and Phos-tag SDS-PAGE (**Fig. 2A**, bottom) to assess the relative level of phosphorylation. Intact mass spectrometry was employed to quantify phosphorylation levels which revealed variants with an average of 4 (Orc1^IDR-4P^), 9 (Orc1^IDR-9P^), or 11 (Orc1^IDR-11P^) phosphates (**Fig. 2B**).

**Figure 2:**
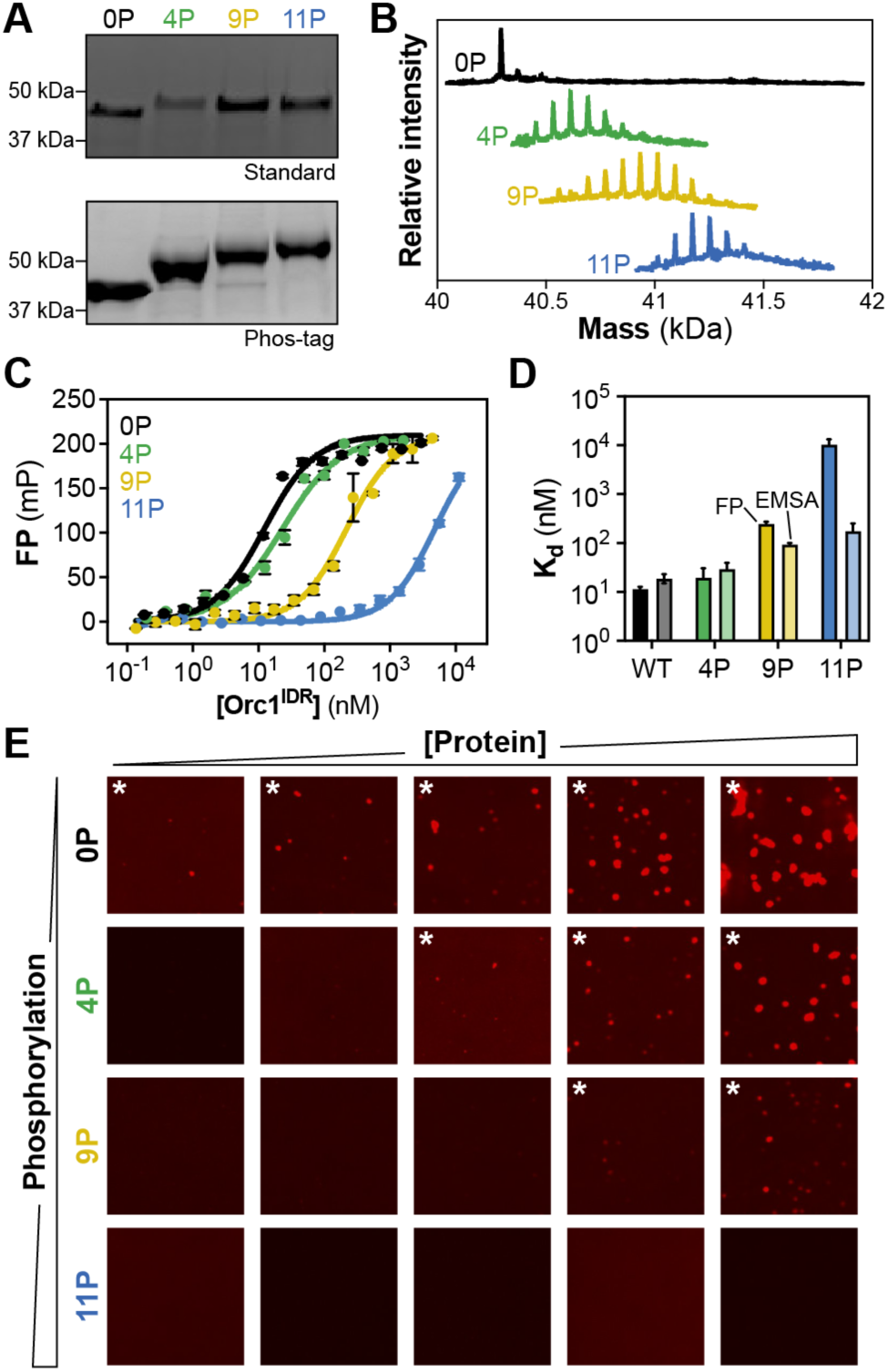
Phosphorylation tunes Orc1 IDR function. A, analysis of Orc1 IDR phospho-variants by normal (top) and Phos-tag (bottom) SDS-PAGE. Gels are Coomassie stained. B, intact mass spectrometry analysis of Orc1 IDR phospho-variants. C, the DNA binding affinity of Orc1 IDR phospho-variants was measured by fluorescence polarization (FP). D, summary of dissociation constants (K_d_) measured by fluorescence polarization (darker shade) or EMSAs (lighter shade) for Orc1 IDR phospho-variants. E, phase separation analysis of Orc1 IDR phospho-variants. Each phospho-variant was titrated from 125 nM to 2 µM (left to right with 2-fold dilutions between) with stoichiometric amounts of Cy5-labeled duplex DNA and imaged by confocal fluorescence microscopy.

To test the mechanism of phospho-regulation, we compared the DNA binding affinity and phase separation propensity of the purified Orc1^IDR^ phospho-variants. The *Drosophila* Orc1 IDR binds DNA non-specifically (9) and in all experiments a fluorescently labeled 60 base pair duplex DNA with approximately 50% GC content was used as a substrate (see **EXPERIMENTAL PROCEDURES** for DNA sequence). Fluorescent polarization (FP) (**Fig. 2C**) and electrophoretic mobility shift assays (EMSAs) (summary data in **Fig. 2D** and raw data in **Fig. S1A-D**) were used to measure DNA binding affinity. Consistent with prior results (10), FP measurements revealed low nanomolar DNA binding affinity for the non-phosphorylated variant (**Fig. 2C**, black line, K_d_ = 12.4 nM) and we observed progressively weaker DNA binding affinity as the level of phosphorylation was increased (Orc1^IDR-4P^ K_d_ = 23.5 nM, Orc1^IDR-9P^, K_d_ = 222 nM, and Orc1^IDR-11P^ K_d_ ≈ 5 µM). Overall, EMSAs showed the same trend (**Fig. 2D**) although calculated K_d_’s were generally lower (higher affinity binding) which we suspect is due to sample dilution into running buffer that results in reduced ionic strength. These data show that progressively higher levels of phosphorylation result in progressively weaker DNA binding. This result is consistent with phosphorylation exerting rheostat control on the Orc1 IDR rather than the all-or-none binding expected for switch-like control of differentially phosphorylated variants.

We next assessed whether the phosphorylation-induced loss of DNA binding resulted in a commensurate loss in phase separation propensity (**Fig. 2E**). Consistent with our previous observations (9), we observed phase separation of the non-phosphorylated variant, Orc1^IDR-0P^, down to low nanomolar concentrations (125 nM) (**Fig. 2E**, top row). Conversely, phospho-variants with increasingly higher levels of phosphorylation had increasingly higher critical concentrations (500 nM for Orc1^IDR-4P^ and 1 µM for Orc1^IDR-9P^), and a complete loss of phase separation was observed for the most highly phosphorylated variant, Orc1^IDR-11P^. Although phosphorylation has a graded, rheostat-like effect on Orc1’s DNA binding affinity and critical concentration for phase separation, phosphorylation may nonetheless switch the phase separation propensity of Orc1 on and off if cellular protein levels are set near the protein’s critical concentration. This highlights a potentially interesting aspect of phospho-regulation, where partial phosphorylation of the Orc1 IDR may only marginally impact cellular chromatin binding while fully ablating condensate formation.

### Phosphorylation alters the bulk chemical properties of the Orc1 IDR to tune chromatin binding and cellular patterning

The observed rheostat-like control of Orc1^IDR^ suggests that it is not a particular phosphorylation site(s) that underlies regulation, but rather the cumulative impact that phosphorylation has on the IDRs bulk chemical properties. Indeed, full phosphorylation of the Orc1 IDR (**Fig. 3A**, 15 CDK sites) would lower the protein’s net charge from +23.8 to −2.6 (each phosphate bears an ≈ −1.75 charge at pH 7.5), a dramatic chemical change (**Fig. 3B**). If regulation results simply from changing the overall charge of the sequence, then a phospho-mimetic mutant, such as by introduction of an aspartate/glutamate at phospho-sites, would impact Orc1^IDR^ function only to the extent that it changes the total net charge of the protein. Since aspartate/glutamate (z = −1) has a lower charge density than a phosphate ion (z = −1.75), a phospho-mimetic mutant of the fully phosphorylated state would introduce only 15 negative charges (**Fig. 3B**, white symbols), lowering the net charge to z = 8.8 (**Fig. 3B**). We therefore produced a phospho-mimetic mutant of the Orc1 IDR (Orc1^IDR-PM^, “[S/T]P” → “DP”, **Fig. 3A**) and, as a test of how bulk chemical properties alter Orc1^IDR^ function, compared its DNA binding and phase separation propensity to Orc1^IDR^ phospho-variants.

**Figure 3:**
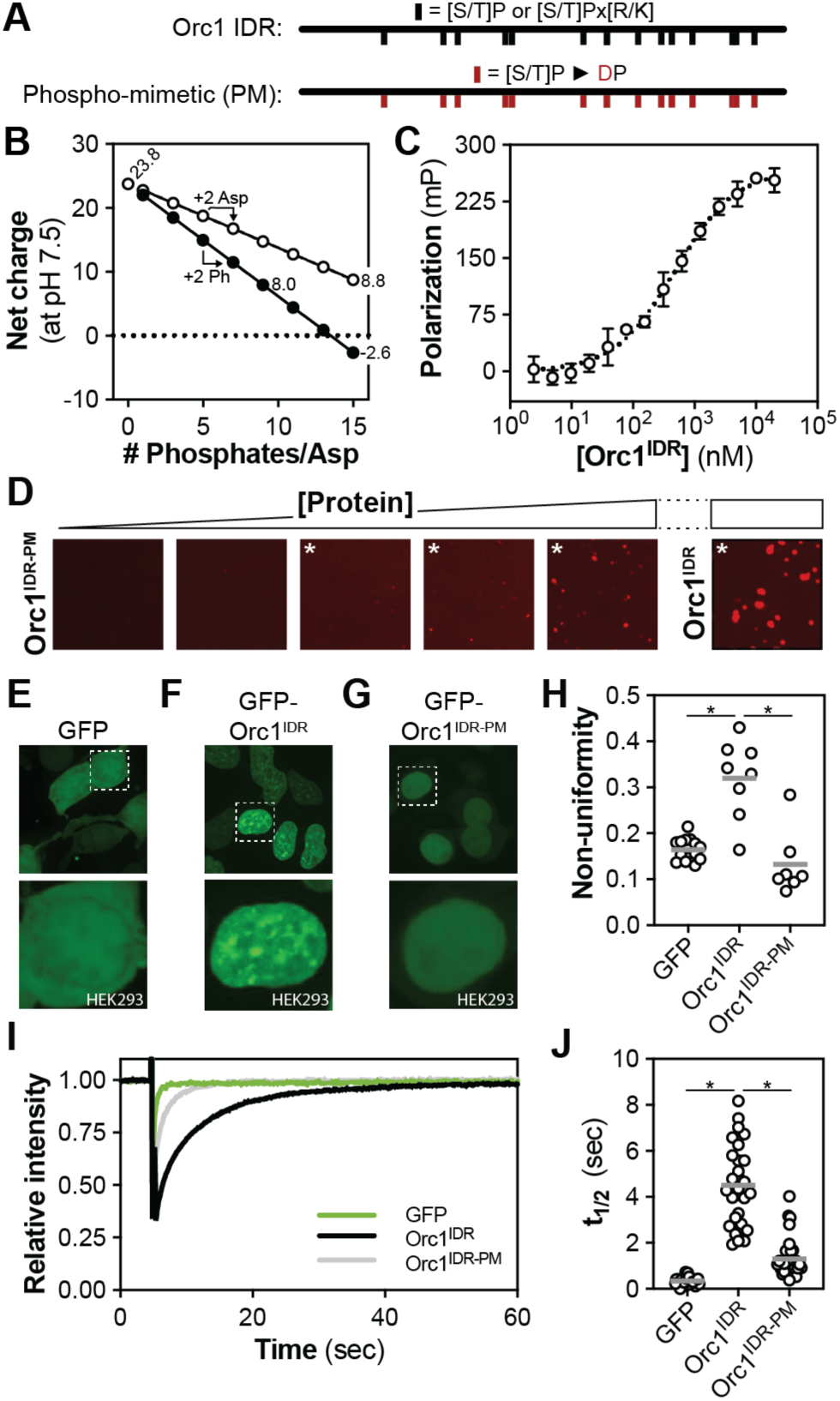
Orc1 IDR phosphorylation regulates chromatin binding and heterochromatin partitioning. A, position of phosphorylation (black hash marks) or phospho-mimetics (red hash marks) in wild-type Orc1 IDR (top, Orc1^IDR^) or the phospho-mimetic mutant (bottom, Orc1^IDR-PM^). B, change in Orc1^IDR^ net charge with progressive phosphorylation (black-filled markers) or introduction of phospho-mimetics (white-filled markers). C, the DNA binding affinity of Orc1^IDR-PM^ was measured by fluorescence polarization (FP). D, phase separation analysis of Orc1^IDR-PM^. Orc1^IDR-PM^ was titrated from 125 nM to 2 µM (left to right with 2-fold dilutions between) with stoichiometric amounts of Cy5-labeled duplex DNA and imaged by confocal fluorescence microscopy. E-G, cellular distribution (50 µm x 50 µm top panel) and zoom view of nuclear distribution (15 µm x 15 µm bottom panel) of E, GFP, F, GFP-tagged Orc1^IDR^, or G, GFP-tagged Orc1^IDR-PM^ in HEK293 cells. H, non-uniformity of nuclear GFP intensity in HEK293 cells expressing constructs in E-G. Each marker is a single cell non-uniformity score, and the population mean is indicated by the grey horizontal line. I, analysis of chromatin binding in HEK293 cells using FRAP for GFP (green line), GFP-Orc1^IDR^ (black line) and GFP-Orc1^IDR-PM^ (grey line). J, the half time of recovery (t_1/2_) was calculated from FRAP curves (I). Each marker represents the t_1/2_ calculated for a single FRAP experiment in a single cell and the population mean is indicated by the grey horizontal line.

Using FP (**Fig. 3C**) and EMSA (**Fig. S2A**) we assayed the DNA-binding affinity of Orc1^IDR-PM^ and compared it to the range of affinities observed for our phospho-variants. EMSA analysis revealed a qualitative loss in Orc1^IDR-PM^’s DNA-binding affinity compared to Orc1^IDR-0P^ (**Fig. S2A**) and fluorescence polarization measurements showed that Orc1^IDR-PM^ binds DNA with modest affinity (**Fig. 3C**, K_d_ = 546 ± 54 nM). Consistent with our predictions, the DNA binding affinity of Orc1^IDR-PM^ is markedly reduced compared to the unphosphorylated Orc1 IDR (**Fig. 2C**, black line, K_d_ = 12.4 nM) but retains tighter binding than Orc1^IDR-11P^ (**Fig. 2C**, blue line, K_d_ ≈ 5 µM), a variant with fewer modified groups but an overall lower net charge. We also assessed the phase separation propensity of Orc1^IDR-PM^. We prepared a one-dimensional phase separation screen by varying the concentration of Orc1^IDR-PM^ from 125 nM up to 2 µM and, after adding stoichiometric amounts of DNA, we imaged the reactions by fluorescence microscopy. Compared to non-phosphorylated Orc1^IDR^, which phase separates down to 125 nM, we observed a significant increase in the critical concentration of Orc1^IDR-PM^ to 500 nM (**Fig. 3D**). These data support a model where phosphorylation tunes DNA binding and phase separation by altering Orc1^IDR^’s bulk chemical properties.

The identification of an Orc1 mutant with weakened DNA binding and phase separation propensity provided the opportunity to test the impact of phase separation on Orc1 spatial patterning in cells. We therefore expressed enhanced Green Fluorescence Protein (GFP)-tagged Orc1^IDR^ and Orc1^IDR-PM^ in HEK293 cells and assessed their spatial distribution using spinning disk confocal fluorescence microscopy (**Fig. 3E-G**). As a negative control, GFP alone was expressed and imaged (**Fig. 3E**). As observed previously for full-length Orc1 (23, 27), Orc1^IDR^ showed a punctate distribution in interphase nuclei (**Fig. 3F**). Prior work shows that the non-uniform nucleoplasmic patterning represents the enrichment of ORC within heterochromatin (14, 16), which we confirmed by imaging cells co-expressing fluorescently tagged Orc1^IDR^ and human Hp1*a* (**Fig. S2B**). On the other hand, Orc1^IDR-PM^ was uniformly distributed throughout the nucleus (**Fig. 3G**) and did not form foci, similar to GFP’s nuclear distribution pattern (**Fig. 3E**). To quantify the effect on foci formation, we calculated the non-uniformity of GFP signal in the nucleus and observed a robust reduction in the non-uniformity of Orc1^IDR-PM^ compared to Orc1^IDR^, with the phospho-mimetic IDR and GFP being similarly uniform (**Fig. 3H**). We used the same constructs in fluorescence recovery after photobleaching (FRAP) experiments to determine whether the reduced DNA-binding affinity of Orc1^IDR-PM^ translates to weaker chromatin binding in cells (**Fig. 3I-J**). FRAP curves were collected from more than twenty cells for each construct and were fit with an exponential one-phase association to calculate the halftime of recovery (t_1/2_, **Fig. 3J**). Consistent with its known chromatin binding capabilities (10), Orc1^IDR^ (**Fig. 3I**, black curve, t_1/2_ = 4.5 sec) showed significantly slower recovery after bleaching than GFP (**Fig. 3I**, green curve, t_1/2_ = 0.3 sec). Conversely, recovery of Orc1^IDR-PM^ (**Fig. 3I**, grey curve, t_1/2_ = 1.3 sec) was significantly faster and indicates that this variant has reduced chromatin binding affinity, consistent with the phosphorylation-induced reduction in *in vitro* DNA binding (**Fig. 3C**).

### Regulatory phosphorylation does not depend on specific phospho-sites but instead requires their equitable distribution

Our data demonstrate that phosphorylation regulates Orc1 IDR function simply by reducing the net charge on the protein. This suggests that the precise positioning of phosphorylation sites may not be important. To test this, we first assessed the sequence conservation (**Fig. 4A**) and spatial distribution (**Fig. 4B**) of CDK sites in nine arthropod Orc1 IDR orthologs. Sequence conservation analysis revealed that only a single phosphorylation site is universally conserved across this set of sequences (**Fig. 4A** and **Fig. 4B**, blue). As expected, comparison of more closely related orthologs (e.g., *Bactrocera tryoni* (“Bt”) and *Rhagoletis pomonella* (“Rp”) in **Fig. 4B**) revealed a higher number of positionally conserved CDK consensus sites, although even within these sequence pairs CDK sites appear to be readily gained and lost (**Fig. 4C**). These analyses indicate that lineage-specific changes to CDK site position are well tolerated, and that iterative alterations over long periods of time (241 million years in the **Fig. 4B** phylogenetic tree) have shuffled phospho-site position while preserving both the overall density of sites (**Fig. 1**) and their relatively equitable distribution across the length of the sequence (**Fig. 4B**).

**Figure 4:**
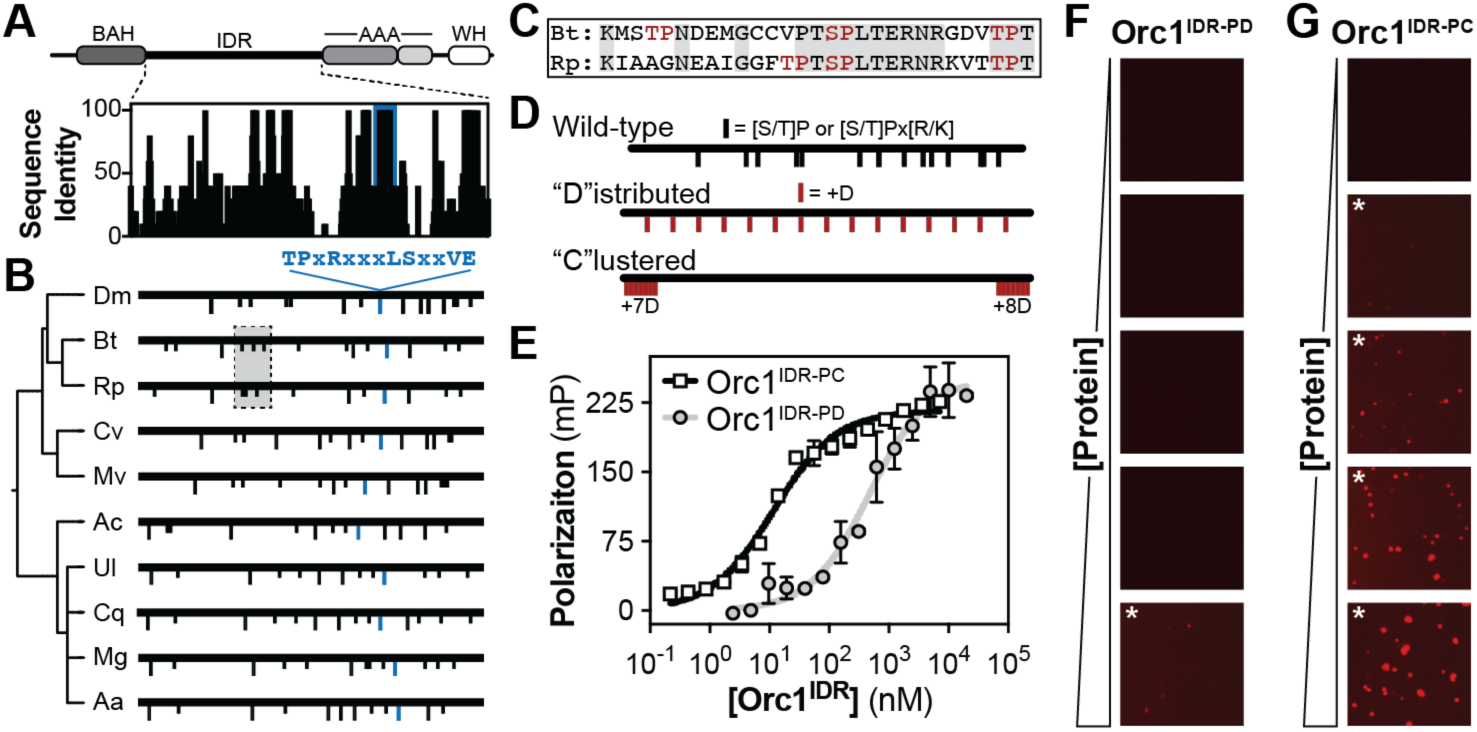
Phospho-regulation of Orc1^IDR^ does not depend on specific phosphorylation sites. A, sequence identity plot for the Orc1 IDR calculated from the alignment of nine arthropod orthologs. B, position of full (long hash mark) and minimal (short hash mark) CDK phosphorylation motifs in nine arthropod Orc1 IDR orthologs. C, pairwise sequence alignment of grey-shaded region in panel B revealing shuffling of CDK phosphorylation motifs. D, position of phosphorylation (black hash marks) and phospho-mimetics (red hash marks) in wild-type Orc1 IDR (Orc1^IDR^) compared to the phospho-distributed (Orc1^IDR-PD^) and phospho-clustered mutants (Orc1^IDR-PC^). E, the DNA binding affinity of Orc1^IDR-PD^ and Orc1^IDR-PC^ was measured by fluorescence polarization (FP). F-G, phase separation analysis of F, Orc1^IDR-PD^ and G, Orc1^IDR-PC^. Each protein was titrated from 125 nM to 2 µM (top to bottom with 2-fold dilutions between) with stoichiometric amounts of Cy5-labeled duplex DNA and imaged by confocal fluorescence microscopy.

We next designed phospho-mimetic variants to experimentally test whether the precise position of phosphorylation sites is important for Orc1 IDR regulation. Specifically, we produced two variants that contained fifteen phospho-mimetic aspartates that were inserted either regularly throughout the sequence (“P”hospho-“D”istributed, Orc1^IDR-PD^) or irregularly into an N and C-terminal cluster (“P”hospho-“C”lustered, Orc1^IDR-PC^) (**Fig. 4D**). The regular spacing of phospho-mimetic residues in Orc1^IDR-PD^ is similar to the relatively equitable distribution of CDK motifs in the native protein. In neither variant do the positions of phospho-mimetic residues overlap with any native CDK motif. By comparing the DNA binding (**Fig. 4E**) and phase separation propensity (**Fig. 4F-G**) of these two variants we tested the importance of phospho-site position versus distribution in regulating Orc1 IDR function. Fluorescence polarization DNA binding assays revealed that Orc1^IDR-PC^ retained wild-type like DNA binding affinity (**Fig. 4E**, black line, K_d_ = 12 ± 1 nM) but that Orc1^IDR-PD^ had significantly reduced affinity (**Fig. 4E**, grey line, K_d_ = 590 ± 197 nM), comparable to that of Orc1^IDR-PM^ (**Fig. 3C**, K_d_ = 546 nM), the phospho-mimetic variant with aspartates positioned at the native CDK sites. Finally, we assayed the phase separation propensity of these two variants (**Fig. 4F-G**). Consistent with our DNA binding analysis, we observe a major reduction in the phase separation propensity of Orc1^IDR-PD^ (**Fig. 4F**, critical concentration = 2 µM) compared to the non-phosphorylated Orc1^IDR^ (**Fig. 2E**, critical concentration = 125 nM), and only a marginal change in Orc1^IDR-PC^ (**Fig. 4G**, critical concentration = 250 nM). These data demonstrate that phospho-regulation of the Orc1 IDR does not require phosphorylation at specific sites but does depend on the sites being well-distributed throughout the sequence.

### Cellular regulation of Orc1 does not require specific phosphorylation sites

We next asked if phospho-regulation of Orc1 cellular patterning is as insensitive to phospho-site position as our *in vitro* studies suggest. We therefore expressed both the phospho-distributed (Orc1^IDR-PD^) and phospho-clustered (Orc1^IDR-PC^) Orc1 IDR mutants in HEK293 cells. In addition to the fifteen aspartate insertions, these cellular constructs also contained mutations to all fifteen native phospho-sites (“[S/T]P” to “AP”) to prevent hyper-phosphorylation of the protein. Like the Orc1 IDR phospho-mimetic (Orc1^IDR-PM^, **Fig. 3G**), Orc1^IDR-PD^ was uniformly distributed and did not form heterochromatic puncta (**Fig. 5A**). Conversely, Orc1^IDR-PC^ formed nuclear foci (**Fig. 5B**), similar to the behavior of the wild-type sequence (**Fig. 3F**). We quantified this behavior by calculating nuclear non-uniformity and observed significantly higher non-uniformity for Orc1^IDR-PC^ compared to Orc1^IDR-PD^ (**Fig. 5C**), although the non-uniformity of the clustered variant was still less than what is observed for the wild-type protein. We additionally compared the chromatin binding activity of these two variants using FRAP (**Fig. 5D-E**). Consistent with our *in vitro* DNA binding results (**Fig. 4E**), Orc1^IDR-PC^ had a relatively slow half-time of recovery that is consistent with chromatin binding (**Fig. 5D**, black line), while Orc1^IDR-PD^ had a significantly faster half-time of recovery (**Fig. 5D**, grey line). The t_1/2_ for Orc1^IDR-PC^ (t_1/2_ = 3.8 sec) and Orc1^IDR-PD^ (t_1/2_ = 1.5 sec) (**Fig. 5E**) is similar to that of Orc1^IDR^ (t_1/2_ = 4.5 sec) and Orc1^IDR-PM^ (t_1/2_ = 1.3 sec) (**Fig. 3J**), respectively. These data demonstrate that phospho-regulation of Orc1 cellular patterning and chromatin binding depends only on the equitable distribution of Orc1 IDR phospho-sites and not on their precise position in the sequence.

**Figure 5:**
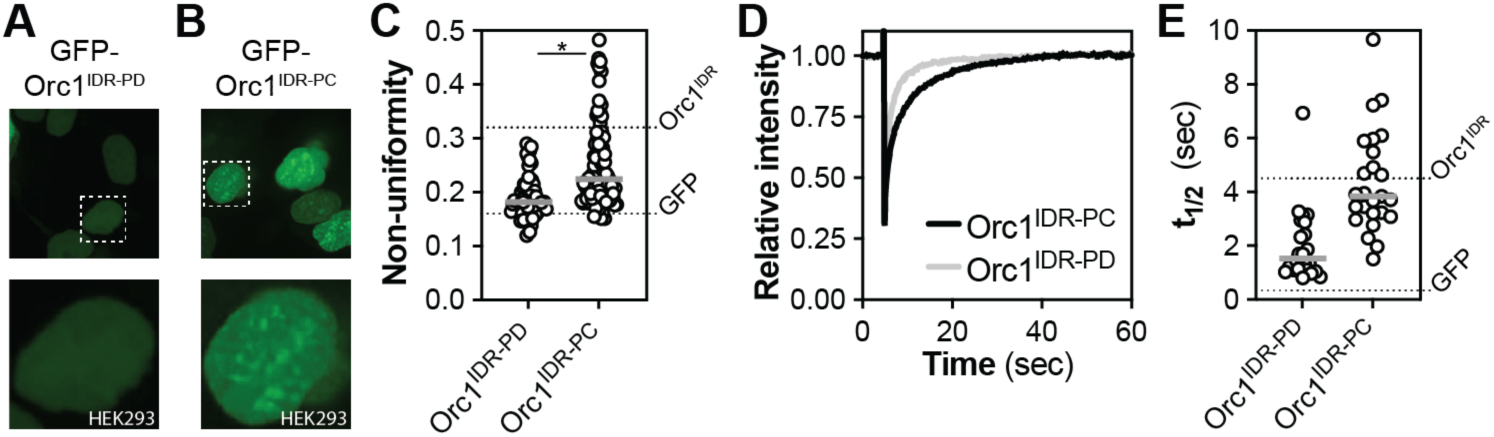
Regulation of Orc1 IDR cellular patterning is insensitive to phospho-site position. A-B, cellular distribution (50 µm x 50 µm top panel) and zoom view of nuclear distribution (15 µm x 15 µm bottom panel) of A, GFP-tagged Orc1^IDR-PD^ and B, Orc1^IDR-PC^ in HEK293 cells. C, non-uniformity of nuclear GFP intensity for GFP-Orc1^IDR-PD^ and GFP-Orc1^IDR-PC^ expressing HEK293 cells. Each marker is a single cell non-uniformity score, and the population mean is indicated by the grey horizontal line. D, analysis of chromatin binding in HEK293 cells using FRAP for GFP-Orc1^IDR-PD^ (grey line) and GFP-Orc1^IDR-PC^ (black line). E, the half time of recovery (t_1/2_) was calculated from FRAP curves (D). Each marker represents the t_1/2_ calculated for a single FRAP experiment in a single cell; the population mean is indicated by the grey horizontal line.

## DISCUSSION

Here we demonstrate that phosphorylation of the Orc1 intrinsically disordered region regulates recruitment of Orc1 to heterochromatin. Instead of a specific phosphorylation “code”, we find that the functional impact of phosphorylation derives from an overall change to the bulk chemical properties of the Orc1 IDR. Although the precise position of phospho-sites can be altered without a measurable functional impact, we find that the equitable distribution of these sites across the length of the sequence is important and that clustering phospho-mimetics ablates their regulatory effect. Consistently, sequence analyses show that while the high density of Orc1 IDR phospho-sites is conserved in animals, as is their equitable distribution across the sequence, specific sites are readily gained and lost. These studies fill an important gap in our understanding of the molecular interactions that underlie regulated recruitment of Orc1 to heterochromatin.

Heterochromatin is a complex, heterogenous compartment and ORC has multiple heterochromatin-specific interaction partners: Orc1 and Orc3 bind Hp1*α* (14, 16) and Orc2-5 bind ORCA (21). Whether the recruitment of ORC to heterochromatin depends on cooperative or hierarchical interactions within this network is not currently known, but available data suggest that the recruitment of heterochromatin resident factors may be interdependent (16, 28). Our data demonstrate that phosphorylation plays a dominant role in regulating Orc1’s partitioning into heterochromatin. While the effect of phosphorylation could result from regulation of protein-protein interactions, we think this is unlikely for several reasons. First, we observe cross-species compatibility with the fly Orc1 IDR able to target human heterochromatin. This, combined with an absence of sequence similarity between the fly and human Orc1 IDRs (10), suggests that specific protein-protein interactions might be dispensable for heterochromatin recruitment. Second, inhibition of protein-protein interactions would likely depend on phosphorylation at specific sites; however, we observe that evolution has scrambled the position of CDK motifs (**Fig. 4A-B**), and that phospho-site position can be altered whilst maintaining the ability to negatively regulate heterochromatin recruitment (**Fig. 5A**). These observations prompt the consideration of alternative mechanisms to explain phospho-regulated recruitment of Orc1 to heterochromatin.

Certain factors are recruited to heterochromatin through direct interactions with DNA. These include the recently discovered human ZNF512 and ZNF512B (29), and the AT-hook containing human HMGA family of proteins (30) and *Drosophila* D1 (31), which bind heterochromatic DNA with varying levels of specificity. The Orc1 IDR may likewise be recruited to heterochromatin through direct DNA binding.

However, instead of relying on sequence-specific DNA binding (an activity the Orc1 IDR lacks), the Orc1 IDR’s specificity for heterochromatin would derive from the local unveiling of its non-specific DNA binding activity. Notably, protein phosphatase 1 (PP1), which dephosphorylates ORC (including Orc1 and Orc2 subunits) (32–34), forms a stable complex with Rap1-interacting factor 1 (Rif1) (35, 36), and these are together enriched in heterochromatin where they regulate replication timing (37). We thus propose that dephosphorylation and activation of the Orc1 IDR’s DNA binding activity occurs specifically in the vicinity of heterochromatin. In support of this mechanism, prior work shows that depletion of Rif1 or inhibition of PP1 reduces levels of chromatin-associated Orc1 and Orc2 (33, 34) and that Rif1 and PP1 co-immunoprecipitate with Orc1 in human cells (23). Interestingly, although both *Drosophila* and human Orc1 IDRs contain a PP1 docking motif (**Fig. S3A-B**), these motifs are at different positions in the IDR and are also of a different type, with the fly ortholog possessing a “SILK” motif and the human ortholog an “RVxF” motif (38). This demonstrates that despite the marked divergence of Orc1 IDRs, evolution has maintained the presence, but not the precise position or sequence, of short regulatory motifs. Altogether, this highlights a potentially novel mechanism for the recruitment of Orc1, and possibly other chromatin binding proteins, to specific chromosomal loci, requiring a regulatable non-specific DNA binding activity and the enrichment of regulatory factors at specific chromosomal locations.

The mechanism we describe for site-specific chromosome binding could also underlie other aspects of Orc1 function. In the early fly embryo, replication licensing occurs in late mitosis of each cell cycle (there are no gap phases) and origins of replication are closely spaced to facilitate the rapid cell cycles of the early embryo (39–42). Consistently, we and others have observed that ORC is recruited to and uniformly coats mitotic chromosomes specifically in anaphase (43) and that CDK-dependent phosphorylation of the Orc1 IDR is responsible for this temporal regulation (9, 10). Intriguingly, prior work shows that the Repo-Man•PP1 complex, which is required for mitotic chromosome condensation, is recruited to chromatin in early anaphase coincident with Orc1 (44). Akin to site-specific dephosphorylation of Orc1 in heterochromatin by Rif1•PP1, the chromosome-bound Repo-Man•PP1 complex may also facilitate the dephosphorylation of the Orc1 IDR in the vicinity of chromatin to drive mitotic chromosome binding. Such a mechanism would ensure that the Orc1 IDR, which will bind a variety of negatively charged polymers (including RNA and heparin (45)), targets chromatin specifically. Extending this further, it is plausible that phosphatases enriched at specific chromosomal loci could locally regulate the Orc1 IDRs DNA binding activity to facilitate the more stringent origin selection that is seen during interphase in differentiated cells (46).

These studies reveal the mechanism for the regulated recruitment of Orc1 to heterochromatin and, more broadly, suggest a new model for the recruitment of ORC to specific chromosomal loci. Intriguingly, our phylogenetic and biochemical studies demonstrate that this regulation does not depend on any specific phosphorylation site but instead on the equitable distribution of sites throughout the sequence, which change the overall chemistry of the Orc1 IDR to reduce DNA binding and phase separation propensity. These findings motivate future studies to test the contribution of chromatin-bound phosphatases in directing ORC to specific chromosomal sites.

## EXPERIMENTAL PROCEDURES

### Computational studies of phospho-site density

A custom Python script was written to calculate the density of CDK sites proteome wide (**Fig. 1B**). Briefly, we analyzed the entire *Drosophila* proteome (UniProt dataset UP000000803_7227.fasta) using Metapredict to identify all disordered regions longer than fifty amino acids. For each disordered region, we calculated the number of both full (‘[S/T]PX[R/K]’) and minimal (‘[S/T]P’) CDK consensus motifs and normalized the data by the length of each sequence. This CDK site density data was then analyzed in GraphPad Prism as a cumulative distribution function. Likewise, the number (**Fig. 1C**) and position (**Fig. 4B**) of CDK sites in Orc1 IDR orthologs was calculated with a custom script and visualized using GraphPad Prism. To determine whether the prevalence of CDK consensus motifs was influenced by the overall amino acid composition of the sequence (**Fig. 1D**), we wrote a custom Python code to randomly scramble each Orc1 IDR ortholog 10,000 times, count the number of CDK (full and minimal) and Casein Kinase II (‘[S/T]XX[D/E]’) motifs in each randomized sequence, and then calculate the average number of sites in all 10,000 random scrambles. These data were visualized in GraphPad Prism by plotting the averages for each ortholog against the observed number of CDK or Casein Kinase II sites in the wild-type sequences. This code is freely available at our GitHub repository (https://github.com/MWPlabUTSW/Adiji2026.git).

### Cloning, expression, and purification of *Drosophila* CDK2•CycE

The *Drosophila* CDK2•CycE complex was expressed and purified in *Spodoptera frugiperda* (Sf9) insect cells with the aid of the baculovirus expression system as previously described (10) with some minor modifications. Specifically, we re-engineered vector 438C (QB3 MacroLab) to remove the His6 tag and cloned CDK2 into the resulting vector, which we have named 438-MBP, for expression as an N-terminal Maltose Binding Protein (MBP) fusion protein with a TEV protease cleavable tag. *Drosophila* CycE was cloned into vector 438B (QB3 MacroLab) for expression as an N-terminal His6 fusion protein with a TEV protease cleavable tag. These constructs were used to generate recombinant bacmid DNA following previously established protocols (10) and subsequently transfected into Sf9 cells for generation of high titer virus and expression.

Sf9 cells (Expression Systems Cat. #94-001F) were maintained in ESF 921 media (Expression Systems Cat. #96-001-01) at 27°C and bacmid DNA was transfected into Sf9 cells using Cellfectin II following the manufacturer’s instructions (Thermo Fisher Scientific Cat. #10362100) and incubated for 4 - 7 days at 27 °C. The resulting P0 viral stock was harvested and further amplified in two subsequent rounds of infection to generate P1 and P2 viral stocks. For large-scale expression, Sf9 cells (3×10^6^ cell/ml) were co-infected with CDK2 and CycE P2 viral stocks at a 1:50 virus to cell volume ratio. Cells were then incubated for 48 hr with gentle shaking (120 rpm) before harvesting. Cell pellets were harvested by centrifugation at 5,000 rpm for 15 min, the supernatant was discarded, and the cell pellet stored at −80°C until protein purification.

To purify the CDK2•CycE complex, cell pellets from 2 L of culture were used for purification by resuspending in 80 mL of lysis buffer (50 mM Tris-HCl (pH 7.5), 300 mM KCl, 50 mM imidazole, 10% (v/v) glycerol, 200 μM PMSF, 1 mM β-mercaptoethanol (BME), 1 μM benzonase (Millipore Sigma Cat. #9025-65-4), and 1× cOmplete EDTA-free protease inhibitor cocktail (Thermo Fisher Scientific Cat. #A32965). Cells were lysed on ice by sonication using a Branson Digital Sonifier 450 running five cycles of 15 sec at 100% power followed by 1 min on ice. The lysates were then clarified by centrifugation at 18,000 rpm for 1 hr at 4 °C, and the supernatant was filtered through a 0.45 µm aPES membrane (Nalgene Rapid-Flow, Thermo Fisher Cat. #FB12566511). The filtered lysate was subject to an ammonium sulfate precipitation (975 mM) and incubated under gentle rotation for 30 min at 4°C. The lysate was again clarified by centrifugation at 18,000 rpm for 1 hr and the supernatant was further filtered before loading onto a 5 mL HisTrap HP column (Cytiva Cat. #17524802). The column was washed with twelve column volumes (CV) of wash buffer (50 mM Tris-HCl (pH 7.5), 300 mM KCl, 50 mM imidazole, 10% glycerol, 1 mM BME) and then eluted with a linear gradient of 50 to 250 mM imidazole. The nickel elution fraction was subsequently purified over amylose resin (New England Biolabs Cat. # E8021L). After loading the protein, the resin was washed with three CV of amylose wash buffer (50 mM Tris-HCl (pH 7.5), 300 mM KCl, 10% glycerol, 1 mM BME) and then eluted with two CV of amylose elution buffer (50 mM Tris-HCl (pH 7.5), 300 mM KCl, 10% glycerol, 1 mM BME, 20 mM maltose). The eluted CDK2•CycE complex was concentrated using Amicon Ultra-15 centrifugal filters (Thermo Fisher Cat. # 88528), analyzed for purity by SDS-PAGE and Coomassie staining, and the purified protein was finally aliquoted, flash froze in liquid nitrogen, and stored at −80°C.

### Cloning, expression, and purification of Orc1 IDR variants

The coding sequence for the *Drosophila* Orc1 IDR (Orc1^IDR^, residues 187–549), phospho-mimetic variant (Orc1^IDR-PM^, Ser/Thr to Asp mutations at all native CDK sites), phospho-distributed variant (Orc1^IDR-PD^, fifteen additional aspartates were distributed equitably across the IDR sequence), phospho-clustered variant (Orc1^IDR-PC^, fifteen additional aspartates were added, seven at the N-terminus and eight at the C-terminus), and phospho-dead variant (Orc1^IDR-PDead^, Ser/Thr to Ala mutations at all native CDK sites) were obtained by either gene synthesis (Twist Biosciences) or PCR (see **Supplementary File 1** for list of sequences). Each variant was inserted into vector 1C (QB3 Macrolab) using ligation independent cloning (LIC) for expression in *Escherichia coli* BL21(DE3) cells as a TEV-cleavable N-terminal His6-MBP fusion protein. For expression, overnight cultures were used to inoculate large-scale cultures (800 mL) grown at 37°C with shaking at 250 rpm. After growth to an optical density (OD600) = 0.8, the culture was placed in an ice bath for 15 min and then expression induced by addition of isopropyl β-D-1-thiogalactopyranoside (IPTG) to a final concentration of 1 mM. The induced cells were grown at 20°C for 18 hr with shaking (200 rpm) and then harvested by centrifugation at 5,500 rpm for 15 min at 4°C. The cell pellets were collected and stored at −80°C until purification.

Proteins were purified from the cell pellets of 2 L of culture. The cell pellets were resuspended in 80 mL of lysis buffer (20 mM Tris-HCl, pH 7.5, 500 mM NaCl, 30 mM imidazole, 10% glycerol, 200 µM PMSF, 1× cOmplete EDTA-free Protease Inhibitor Cocktail (Thermo Fisher Scientific Cat. #A32965), 1 mM β-mercaptoethanol (BME), and 0.1 mg/ml lysozyme) and lysed by sonication on ice (5 cycles of 15 sec bursts at 100% power followed by 1 min on ice). Lysates were clarified by centrifugation at 18,000 rpm for 1 hr at 4°C, and the supernatants subsequently filtered through a 0.45 µm aPES bottle-top filter unit (Nalgene Rapid-Flow, Thermo Fisher Cat. #FB12566511) before loading onto a 5 ml HisTrap HP column (Cytiva Cat. #17524802) pre-equilibrated with lysis buffer. The column was then washed with 20 column volumes (CV) of nickel wash buffer (20 mM Tris-HCl, pH 7.5, 500 mM NaCl, 30 mM imidazole, 10% glycerol, 200 µM PMSF, 1 mM BME) and eluted with five CV of nickel elution buffer (20 mM Tris-HCl, pH 7.5, 150 mM NaCl, 500 mM imidazole). The sample was further purified by heparin affinity chromatography on a HiTrap Heparin HP column (Cytiva Cat. #17040701). Specifically, the protein was loaded onto the column and then washed with 10 CV of heparin binding buffer (20 mM Tris-HCl, pH 7.5, 150 mM NaCl, 10% glycerol, 1 mM BME, 400 µM PMSF) before eluting with a linear salt gradient (150 mM to 1 M NaCl over ten CV). Subsequently, the His6-MBP tag was removed by overnight digestion with TEV protease (1:20 w/w ratio) at 4°C followed by an ortho nickel step to separate the cleaved protein from the tag. Protein was then concentrated to <1 ml using a 10K Amicon Ultra-15 centrifugal concentrators (Thermo Fisher Cat. #88528) and subject to a final size exclusion chromatography polishing step over a HiPrep 16/60 Sephacryl S-300 HR or Enrich 650 column equilibrated and run in sizing buffer (50 mM HEPES, pH 7.5, 300 mM potassium glutamate, 10% glycerol, 1 mM BME). Purified proteins were then concentrated, aliquoted, flash froze in liquid nitrogen, and stored at −80°C until use.

### Purification and analysis of Orc1 IDR phospho-variants

Large-scale phosphorylation reactions were used to generate phosphorylated variants of the *Drosophila* Orc1 IDR. Specifically, peak size exclusion chromatography fractions of Orc1^IDR^ were treated with a 5:1 molar ratio of Orc1^IDR^ to CDK2•CycE and ATP (4 mM) and incubated at 25 °C for 30 min in sizing buffer (50 mM HEPES, pH 7.5, 300 mM potassium glutamate, 10% glycerol, 1 mM BME). To obtain higher levels of phosphorylation, reactions were further supplemented with a 20:1 molar ratio of Orc1^IDR^ to CDK2•CycE and incubated for additional 30 min. After the phosphorylation reaction was complete, the final reaction mixture was subjected to a final size exclusion chromatography step to remove CDK2•CycE and unincorporated ATP. Peak fractions containing the phosphorylated Orc1^IDR^ were analyzed by SDS-PAGE, pooled, concentrated, aliquoted, flash-froze in liquid nitrogen and stored at −80°C until use. To better discriminate differentially phosphorylated variants, phospho-variants were analyzed by Phos-tag gel electrophoresis (FUJIFILM Biosciences, Cat. #195-17991) (47) and intact mass spectrometry was performed as previously described (10) to obtain a quantitative understanding of the number of phosphates added to each protein.

### Analysis of DNA binding by electrophoretic mobility shift assays (EMSAs) and fluorescence polarization (FP)

For EMSA analysis of DNA binding, serial dilutions of Orc1 IDR variants were prepared in assay buffer (50 mM HEPES (pH 7.5), 150 mM potassium glutamate, 10% (v/v) glycerol, and 1 mM β-mercapto ethanol) and Cy5-labeled double-stranded DNA (5′-Cy5-GAAGCTAGACTTAGGTGTCATATTGAACCTACTATGCCGAACTAGTTACGAGCTATAACC-3′) was then added to a final concentration of 2 nM. The mixtures were incubated at room temperature for 20 min before loading and running samples on a 1% agarose gel at 100 V for 30 min. A ChemiDoc MP Imaging System (Bio-Rad) was used to image gels with filter settings for Cy5. To quantify binding, free DNA band intensities were measured and used to calculate the fraction of DNA bound for each protein concentration. GraphPad Prism was used to plot the fraction DNA bound versus protein concentration and the resulting curves were fit with a one site specific binding with Hill Slope model to calculate dissociation constants (Kd). For the quantified data, each data point represents the mean ± the standard deviation of three experimental replicates.

For FP analysis of DNA binding, serial dilutions of Orc1 IDR variants were prepared in assay buffer (50 mM HEPES (pH 7.5), 150 mM potassium glutamate, 10% (v/v) glycerol, and 1 mM β-mercapto ethanol) and FITC-labeled double-stranded DNA (sequence: 5′-FITC-GAAGCTAGACTTAGGTGTCATATTGAACCTACTATGCCGAACTAGTTACGAGCTATAACC-3′) was then added to a final concentration of 3 nM. The mixtures were incubated at room temperature for 30 min and then 15 µL of each reaction was transferred to a 384-well plate (Greiner Bio-One, Cat. #3820). Fluorescence polarization measurements were made using a CLARIOstar plate reader (BMG LABTECH). The fluorescence polarization values were blanked and corrected for background using a buffer only and DNA only control, respectively. Measurements were taken for three technical and biological replicates, and the data were plotted in GraphPad Prism and fit in the same way as the EMSA data described above.

### Confocal microscopy phase separation (LLPS) assays

To assess liquid-liquid phase separation, each Orc1 IDR variant was mixed with stoichiometric amounts of a Cy3-labeled double-stranded DNA (sequence: 5’-Cy3– GAAGCTAGACTTAGGTGTCATATTGAACCTACTATGCCGAACTAGTTACGAGCTATAAAC-3’) in a total of 20 µL and at the concentrations indicated in the figure and figure legends. The reaction mixture was incubated at room temperature for a minimum of 5 minutes prior to imaging to allow sufficient time for condensate formation. Condensate formation was visualized by spinning disk confocal fluorescence microscopy using a Nikon Ti2E equipped with a Yokogawa CSU X1 spinning disc. A 561 nm laser was used to excite the Cy3 labeled DNA and imaging was done with a 60x oil immersion objective and appropriate filter sets. Images were analyzed using NIS Elements and FIJI.

### Transfection and imaging of Orc1 IDR variants in HEK293T cells

To visualize the cellular patterning of Orc1 IDR variants in HEK293 cells, each variant was cloned into vector 6D (QB3 Macrolab) for expression with a C-terminal GFP tag. We modified Orc1^IDR-PD^ and Orc1^IDR-PC^ such that in addition to the fifteen aspartate insertions, all fifteen native phospho-sites were mutated to alanine (“[S/T]P” were mutated to “AP”) to avoid hyper-phosphorylation of the protein in the cell. Cloning was done using ligation-independent cloning (LIC) according to established methods (QB3 MacroLab) and sequences were verified by either whole plasmid sequencing or Sanger sequencing.

Human embryonic kidney cells (HEK293T, a gift from the lab of Dr. Xiaochen Bai) were maintained in Dulbecco’s modified Eagle medium (DMEM) supplemented with 5% fetal bovine serum and incubated at 37°C and 5% CO₂. For transfection, cells were seeded in 24-well µ-Plates (Ibidi, Cat. #82426) at a density of 2×10⁶ cells/well and then incubated for 24 hr to reach 60-80% confluence. jetPRIME transfection reagent (Polyplus, Cat. #101000046) was used according to the manufacturer’s instructions. Briefly, transfections were performed by diluting 0.5 µg of plasmid DNA into 50 µL jetPRIME buffer, mixed with 1 µL transfection reagent, incubated for 10 min at room temperature, and then added dropwise to the cultures. The plates were gently swirled and incubated at 37°C for 24-72 hr prior to imaging. Subsequently, the transfected cells were moved directly onto the microscope stage and imaged using spinning-disk confocal fluorescence microscopy on a Nikon Ti2E outfitted with a Yokogawa CSU X1 spinning disk. Z-stacks were collected (8 µm depth at 0.3 µm intervals) using 488 nm laser excitation (5-10% laser power), a 60x oil-immersion objective, and 200 ms exposure time.

### Analysis of Orc1 IDR chromatin binding by fluorescence recovery after photobleaching (FRAP)

The chromatin binding propensity of GFP-tagged Orc1 IDR variants was assessed by fluorescence recovery after photobleaching (FRAP) experiments in HEK293T cells using a Nikon Ti2E outfitted with a Yokogawa CSU X1 spinning-disk confocal fluorescence microscope. For each construct, we assessed binding by FRAP in at least 10 cells. For each cell, a small region (approximately 4 µm^2^) was defined within the cell nucleus and was bleached with a 405 nm laser (25% power with 100 µs dwell time). Pre-bleach images were taken, and, after bleaching, the cell was continuously imaged for 1 minute to assess fluorescence recovery within the bleached area. A 488 nm laser (25% power and 100 µs exposure time) and appropriate filter sets were used for imaging GFP-tagged proteins. Image series were analyzed in FIJI using a custom script (**Supplemental File 2**) to subtract background and quantify the fluorescence intensity both within and outside of the bleached area. The fluorescence intensity outside of the bleached area was used to correct for photobleaching and all intensities were normalized to the intensity at time = 0. The normalized fluorescence intensity within the bleached area was plotted in GraphPad Prism as a function of time and fit with a one-phase association nonlinear regression to calculate the half-time of recovery (t_1/2_). The reported FRAP recovery curves are averaged from all individual FRAP experiments.

### Analysis of Orc1 IDR cellular patterning and quantitation of non-uniformity

The non-uniformity of nuclear fluorescence for GFP-tagged Orc1 variants was used as a quantitative measure of how well Orc1 variants partition into heterochromatin. For these assays, a 60x objective was used to collect large images (≅ 300 µm x 300 µm) of cells expressing different GFP-tagged Orc1 IDR variants. These images were then analyzed with a custom FIJI script (**Supplemental File 3**) to isolate individual cells, identify nuclei, and subsequently measure the mean and maximum intensities within the area. These intensities were used to calculate a custom max-to-mean signal non-uniformity score that is adapted from a common signal uniformity metric used for magnetic resonance imaging (MRI) image analysis (48): 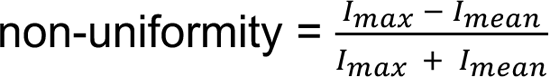, where I_max_ is the maximum pixel intensity and I_mean_ is the mean pixel intensity within the nucleus. The signal non-uniformity was calculated for at least 10 individual cells for each construct, and these values, in addition to their mean, was plotted in GraphPad Prism.

## DATA AVAILABILITY

All raw and analyzed data presented in this paper are publicly available through our lab’s Data Dryad repository (DOI: 10.5061/dryad.1ns1rn988).

## ACKNOWLEDGMENTS

We thank past and present members of the Parker Lab for helpful discussion and technical advice. We also thank Dr. Andrew Lemoff of the UTSW Proteomics Core for technical guidance in assessing phosphorylation by intact mass spectrometry. This work was supported by the National Science Foundation (NSF 2308642, to M.W.P.), the Cancer Prevention and Research Institute of Texas (CPRIT RR200070, to M.W.P.), and the Welch Foundation (V-I-0004-20230731, to M.W.P.). M.W.P is the Cecil H. and Ida Green Endowed Scholar in Biomedical Computational Science.

