## Supplemental Figures for "Phosphorylation alters the bulk chemical properties of Orc1 to tune DNA binding, phase separation, and heterochromatin partitioning"

### SUPPLEMENTAL FIGURES & FIGURE LEGENDS

#### Supplemental Figure 1

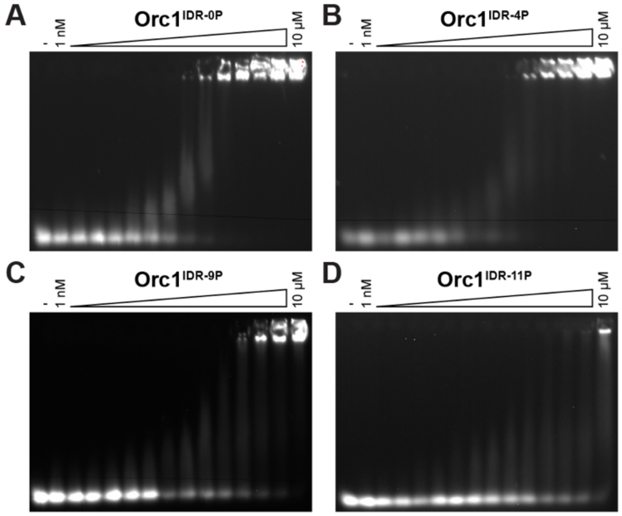


**Supplemental Figure 1: EMSA analysis of phospho-variant DNA binding.** EMSA was used to measure the DNA binding affinity of A, Orc1$^{IDR-0P}$, B, Orc1$^{IDR-4P}$, C, Orc1$^{IDR-9P}$, and D, Orc1$^{IDR-11P}$.

**Supplemental Figure 2**

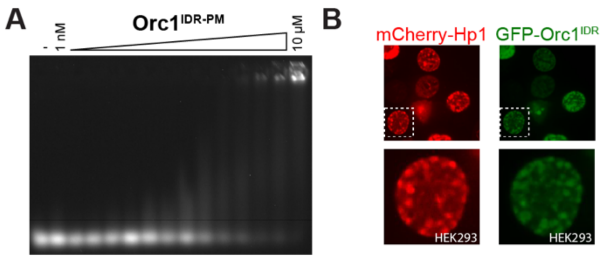


**Supplemental Figure 2: Analysis of the DNA binding affinity and heterochromatin partitioning of the Orc1 IDR phospho-mimetic variant.** A, the DNA binding affinity of Orc1$^{IDR\text{-}PM}$ was measured by EMSA. B, mCherry-tagged human Hp1α (left, red) and GFP-tagged *Drosophila* Orc1$^{IDR}$ (right, green) were co-expressed in HEK293 cells and imaged by confocal fluorescence microscopy. Top panel is a 50 µm x 50 µm field of view and bottom panel is a 15 µm x 15 µm zoom of the cell nucleus outlined by the white dashed line.

**Supplemental Figure 3**

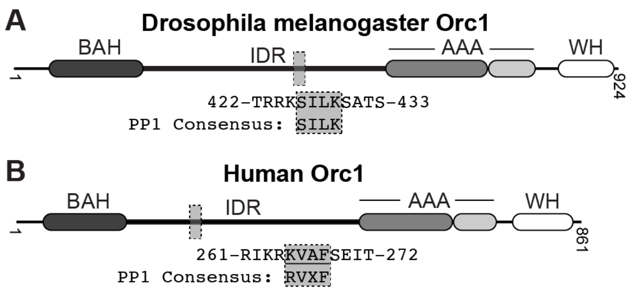


**Supplemental Figure 3: Drosophila and human Orc1 genes have a similar architecture but distinct PP1 docking motifs.** Gene architecture and PP1 docking sites for A, *D. melanogaster* Orc1 and B, human Orc1.

### SUPPLEMENTAL FILES LEGENDS

**Supplemental File 1: Construct sequence file.** This excel file contains the nucleotide and amino acid sequences for all Orc1 IDR variants used in this study.

**Supplemental File 2: FRAP analysis script.** This is a macro written for FIJI to analyze fluorescence recovery after photobleaching (FRAP) data.

**Supplemental File 3: Non-uniformity analysis script.** This is a macro written for FIJI to analyze the nuclear non-uniformity of Orc1 fluorescence signal.
